# Variation in multiple classes of simple sequence repeats can alter drug susceptibility in *Mycobacterium tuberculosis*

**DOI:** 10.64898/2026.09.16.752248

**Authors:** Peter O. Oluoch, Michael J. Luna, Gavin Fujimori, Mayashree Das, Roger Vargas, Kadamba G. Papavinasasundaram, Maha R. Farhat, Christopher M. Sassetti

## Abstract

Insertions and deletions (INDELs) in simple sequence repeats (SSRs) generate relatively high-frequency reversible genetic changes that facilitate bacterial adaptation to changing environments. Analyses of global *Mycobacterium tuberculosis* (*Mtb*) isolates indicate that many SSRs are under diversifying selection, and several of the resulting INDELs in homopolymer tracts (HTs) can increase the pathogen’s fitness during exposure to host and antibiotic stresses. However, the functional impact of most variable SSRs, particularly those within more complex repeat sequences than HT, remains unclear. Here, we combine phylogenomic analysis of clinical *Mtb* strains from Vietnam and Peru with *in vitro* experimental validation of engineered strains to identify SSR INDELs that alter antibiotic susceptibility. Our findings demonstrate that INDELs across multiple SSRs of differing repeat composition are highly variable and correlate with clinical antibiotic resistance. These variants included frameshifting HT INDELs in *ppe13*, *glpK*, *Rv2081c,* and *ppsA*, and in-frame trinucleotide (triplet) SSR INDELS in *ponA1, ppe53,* and *ppe59* that produce much more subtle changes in protein structure. Reconstruction of these INDELs in an isogenic background identified four variants that directly reduce drug potency, including a triplet SSR deletion in *ppe53* that conferred intermediate resistance to isoniazid, rifampicin, and streptomycin. The clinically prevalent *ppe53 CGC_del_* mutation shortens a polyalanine stretch adjacent to the conserved WxG domain and impairs the processing and secretion of the full-length protein. Overall, our work provides additional evidence of selective pressure across *Mtb* SSRs and demonstrates the significance of in-frame INDELs within triplet SSRs, highlighting their contribution to the evolution of antibiotic resistance.

**Author Summary:** Large scale analyses of genome sequences from clinical isolates of *Mycobacterium tuberculosis* have associated many mutations with altered antibiotic susceptibility, informing sequence-based prediction of treatment efficacy. However, previous studies often focus on single nucleotide polymorphisms, limiting our understanding of the roles of more complex mutagenic processes in the adaptation to drug pressure. In particular, insertions and deletions (INDELs) in repetitive regions are quite common, but their impact is incompletely understood. We show that INDELs in simple sequence repeats (SSRs), particularly in-frame INDELs within triplet SSRs, are not only associated with clinical resistance but also confer intermediate antibiotic resistance. In particular, a common one-codon deletion in the *ppe53 gene* shortens a polyalanine stretch in the encoded protein and appears to disrupt critical interactions with the secretion machine, thereby impairing processing and localization of the full-length protein. Our findings reveal a previously unappreciated mechanism of *Mtb* adaptation to antibiotic stress driven by in-frame mutations in triplet SSRs.

## Introduction

Antimicrobial resistance (AMR) remains a major barrier to the successful treatment of tuberculosis (TB), the world’s deadliest infectious disease. According to the 2024 World Health Organization (WHO) Global TB report, a total of 10.7 million people fell ill with the disease, out of which 400,000 cases were multidrug- or rifampicin-resistant (MDR/RR-TB) [1]. Recent advances in whole-genome sequencing (WGS) for clinical isolates of *Mycobacterium tuberculosis* (*Mtb*), the causative agent of TB, have facilitated the characterization of key mutations underpinning antibiotic resistance, culminating in the development of clinically informative catalogs that guide decisions on diagnosis and treatment [2–4]. Beyond canonical high-level resistance conferring mutations in drug targets and prodrug activators, selection for mutations in other genes can produce strains with intermediate resistance, increased antibiotic tolerance, and/or enhanced post-antibiotic exposure recovery [5–9]. Despite the likely importance of these more subtle phenotypes in prolonging therapy, promoting relapse, and facilitating selection for high-level drug resistance [6,10], the identity and impact of individual mutations remain incompletely defined.

Previous genome-wide association studies (GWAS) in *Mtb* have largely focused on single-nucleotide variants (SNVs) and excluded more complex repetitive regions of the genome [11]. More recently, insertions and deletions (INDELS) in simple sequence repeats (SSRs) have emerged as frequent sites of plasticity in the *Mtb* genome [12–15]. SSRs are tandemly repeated short DNA motifs (1–6 bp) that are prone to high-frequency reversible INDEL formation [16]. When present in open reading frames or regulatory elements, INDELs in these repeats can produce phenotypically distinct subpopulations, promoting bacterial survival in changing environments [16–18]. While this process, termed phase variation, is well known to mediate transient adaptation to periodically encountered stresses in other bacteria [18–20], we are only beginning to appreciate its role in the adaptive evolution of *Mtb*. Phase-variable homopolymer tract (HT) INDELs across different genes in pathogenic mycobacteria, including *glpK* [12,13], the *frd* operon [22], and upstream regions of both *espA* and *espR* [14,15], have recently been shown to alter antibiotic susceptibility and adaptation to the host environment. Moreover, large-scale genomic analyses have uncovered a landscape of SSR indels that are under selection during human infection and presumably influence fitness under pressures imposed by antibiotics or the immune system [15]. Despite the abundance of candidate SSR loci, functional validation has remained largely limited to HTs, leaving the broader impact of other SSR indels on *Mtb* antibiotic susceptibility poorly understood.

Expansion and contraction of trinucleotide repeats (henceforth referred to as triplet SSRs) are common in mycobacterial genomes, including *Mtb* [21]. Triplet SSRs encode homopeptide runs that can shape protein structure, protein-protein interactions, and subcellular localization [22,23]. For example, clinically prevalent contractions and expansions within a triplet SSR in the *Mtb ponA1* gene alter the length of a proline homopeptide stretch that mediates interactions with the functionally related peptidoglycan hydrolase, RipA [24,25]. Triplet SSRs are also common in *Mtb* proline-glutamic acid (PE) and proline-proline-glutamic acid (PPE) genes [26], a paralog family that is uniquely expanded in pathogenic mycobacterial genomes. PE/PPE heterodimers have been implicated in outer membrane permeability, nutrient uptake, and antibiotic resistance [27–31]. The association of PE/PPE SNVs with antibiotic resistance implicates variations of this class of proteins in *Mtb* adaptation to antibiotic stress [29–31]. Thus, INDELs within SSR types other than HTs, including triplet SSRs, could play an important role in driving adaptation to antibiotic stress in *Mtb*.

In this study, we integrate *Mtb* phylogenomics with GWAS to define signatures of selection on SSR INDELs and quantify their association with antibiotic resistance. In contrast to previous studies analyzing massive global *Mtb* collections, we focused on distinct, well-curated isolate sets from Vietnam and Peru. This allowed us to account for the specific impacts of local demographics and regional TB control programs on *Mtb* evolution. In addition to HT INDELs, we report that in-frame triplet SSR INDELS, especially those in PE/PPE or membrane/cell wall-associated proteins, are common in Mtb isolates and are significantly correlated with drug resistance. We demonstrate that a highly prevalent *ppe53* triplet SSR INDEL broadly confers intermediate resistance to TB antibiotics by altering the post-translational processing and secretion of the protein. In summary, this work further confirms the clinical and evolutionary significance of SSR INDELs, demonstrating a role for triplet SSR variations in *Mtb* adaptation to antibiotic stress.

## Results

### Selection signatures in SSR INDELs of *M. tuberculosis*

We focused our study on two geographically distinct populations to identify selection signatures that may be specific to different regions. We first curated publicly available whole-genome sequencing datasets of *Mtb* isolates from Vietnam and Peru. Altogether, a total of 1,635 isolates from Ho Chi Minh City (HCMC), Vietnam [32], and 2,106 isolates from Lima, Peru [33,34] were processed (**Fig. 1A**). To detect high-quality insertions and deletions in the repetitive regions of the *Mtb* genome, we applied strict filtering criteria based on sequencing depth, reads mapping rate, variant call quality, empirical base-level recall (EBR) values [35], and variant call concordance after genotype/re-genotyping [36] (*Materials and Methods*). After filtering, 1523 Vietnamese and 1441 Peruvian isolates met the quality threshold for sequencing depth, mapping performance, and call concordance, and successful genotyping in over 75% of the SSR loci (**Fig. 1A***, Supplementary Table 1*). The phylogenetic origins of the *Mtb* isolates included in the downstream analysis reflected the predominant lineages in the two populations [37], lineage 2 (n=984; 64.6%) and lineage 4 (n=1285; 89.1%) in Vietnam and Peru, respectively **(Fig. 1B)**. We excluded SSR INDELs at loci previously shown to have low empirical base-level recall (EBR) values (EBR < 0.9) [35], including all SSR INDELs within genes encoding the PE_PGRS protein family. SNVs in genes known to be associated with *Mtb* drug resistance (DR) [38] were also identified and included in the downstream phylogenetic analysis to serve as controls.

**Figure 1.**
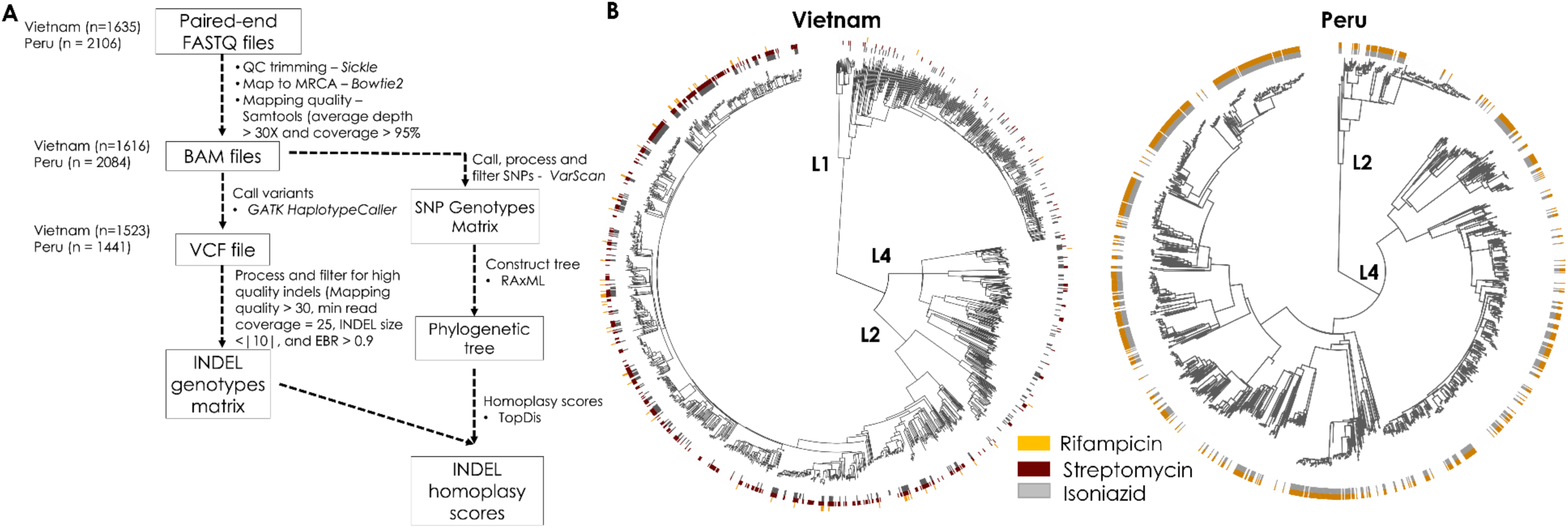
**(A) Bioinformatics workflow of 1,635 (Vietnam) and 2,106 (Peru) publicly available whole-genome sequences.** After the quality control step to exclude low-quality sequences, low sequence depth, and mapping rates, 1616 (Vietnam) and 2084 (Peru) mapped fastq files were used to call repeat indels and SNVs. High-quality SSR indels were retained in 1523 Vietnamese and 1441 Peruvian isolates. A core-SNP-based phylogenetic tree was created using RAxML under the GTR gamma substitution model, and TopDis was used to enumerate the homoplasy scores of each variant. **(B)** Phylogenetic tree of 1523 (Vietnam) and 1441 (Peru) clinical isolates, with phenotypic resistance or sensitivity to isoniazid (grey), streptomycin (red), and rifampicin (orange) marked by the bars.

To estimate the degree of selection on SSR INDELs and DR SNVs across the two populations, we independently calculated homoplasy scores (Hs) using TopDis [15] which quantifies homoplastic events across ancestrally reconstructed phylogenetic trees. Previous simulations based on Mtb mutation rates indicate that an Hs > 5 rarely occurs by chance [15]. Thus, we applied this threshold to our dataset, identifying 243 (Vietnam) and 154 (Peru) SSR INDELs with Hs above this threshold (*Supplementary Table 2*). Known DR-conferring SNVs were among the most homoplastic variants, representing over 25% of mutations with Hs > 50 in both populations. This observation is consistent with the established antibiotic selection signature in these genes [39] and underscores the robustness of the algorithm used. Overall, insertions and deletions within HTs accounted for half of the top 50 variants with the highest Hs in both populations (*Supplementary Table 2*). Among these are variants known to influence *Mtb* fitness. Homoplastic HT INDELs were found in the B11 (Vietnam) and F6 (Peru) sRNAs, both of which are established regulators of mycobacterial virulence [40,41]. Similarly, we identified HT INDELs in the 5’ UTR of *espR*, which modulate adaptation to host and antibiotic-induced stress [14] (**Fig. 2A**).

**Figure 2.**
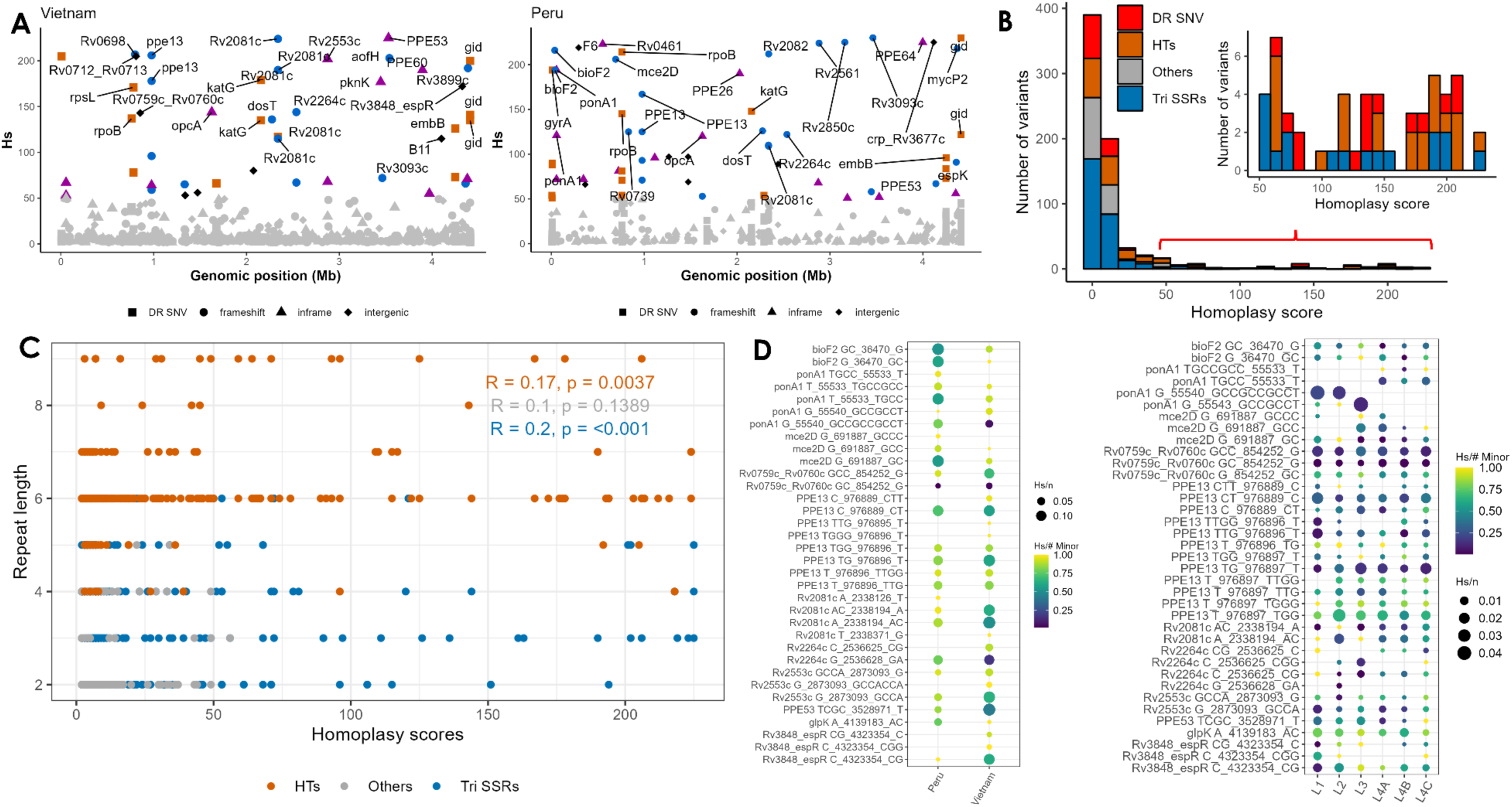
Selection of *M. tuberculosis* simple sequence repeat indels across Vietnamese and Peruvian clinical isolates. **(A)** Homoplasy scores (Hs) of repeat INDELs and DR SNVs across *Mtb* genes. SSR indels are grouped into frameshift, in-frame, or intergenic indels. (**B**) The distribution of homoplasy scores for 745 repeat INDELs in Vietnam, grouped by repeat type (homopolymer tracts, triplet SSRs, or other SSR types). The insert highlights the distribution of the top homoplastic indels across homopolymeric tracts (HTs), triplet repeats (Tri SSRs), and other repeats. (**C**) Relationship between simple sequence repeat length and the homoplasy scores (**D**). Recency ratios (RcR) and homoplasy scores expressed relative to population size for top HTs and Triplet indels across Vietnamese and Peruvian isolates (left) and across common *M. tuberculosis* lineages (right) [15].

A frameshifting pentanucleotide deletion within the GGCGC_3_ SSR of the *Rv3093c* gene was among the top 50 variants in both populations (**Fig. 2A**; *Supplementary Table 2*). *Rv3093c*, which encodes a flavin adenine dinucleotide/flavin mononucleotide (FAD/FMN) reductase, belongs to a gene cluster previously implicated in ethionamide (ETH) resistance [42]. In-frame triplet SSR INDELs were also among the top homoplastic variants in both populations (**Fig. 2B**). These loci included *PPE* family genes (*ppe26*, *ppe53*, *ppe60*, and *ppe64*) and cell-wall/membrane-associated proteins (*ponA1, Rv0461,* and *Rv2553c*) (**Fig. 2A**), providing evidence of ongoing evolution in *Mtb*’s non-frameshifting SSR INDELs. HTs were more variable than other SSR types, but the SSR length, which is a known driver of mutability of these genomic features [43], showed limited correlation with the homoplasy scores across different SSR categories (**Fig. 2C)**. This observation suggests that homoplasy of these INDELs is related to selective pressure rather than frequent mutational events alone.

We next sought to characterize population-specific differences in the selection signatures of the top SSR INDELs. To normalize variation rates across locations and phylogenetic lineages, we estimated the recency ratios (RcR), a measure of recent positive selection expressed as Hs divided by the minor allele count (Hs/#minor) [15], for INDELs across the two geographic populations in our study and across global phylogenetic lineages in a larger published dataset [15]. The majority of the most homoplastic SSR INDELs exhibited an RcR value greater than 0.5 across the two populations and lineages, a signature of more recent evolutionary events. While the majority of SSR INDELs were found across populations and lineages, some exhibited distinct selection patterns (**Fig. 2D)**. For instance, insertion in the *glpK* 7C HT decreases susceptibility to multiple antibiotics [12,13]. This INDEL was significantly more prominent in *Mtb* isolates from Peru than in those from Vietnam (87/1441 in Peru vs 10/1523 in Vietnam, *P-*value < 0.0001, Fisher’s exact test) but did not show strong lineage-specific effects. In contrast, the frequency and selection patterns of INDELs within the *ponA1* CCG triplet SSR varied across *Mtb* lineages. Though annotated as CCG_7_, encoding 7 prolines (7P) in the H37Rv reference genome [44], a fixed CCG > TCG (Pro_631_ > Ser_631_) nonsynonymous mutation is deeply rooted across multiple *Mtb* lineages (**Fig. S1A**). Thus, the ancestral SSR in lineage 4 is CCG_6_TCG (encoding 6 Pro + 1 Ser, 6PS) and CCG_9_TCG (9PS) in lineages 1 and 2 (**Fig. S1B**). This disparity explains the high frequency and low RcRs of the ancestral *ponA1* 9PS (insCCGCCGCCT) variant in lineages 1 and 2 if mapped to the H37Rv reference [15]. In contrast, the *ponA1* 8PS (insCCGCCT) variants in these lineages are infrequent and highly homoplastic, characterized by higher RCR values that indicate more recent evolutionary events (**Fig. 2D**). Within lineage 4 sub-lineages where the 6PS genotype is ancestral (**Fig. S1**), 5PS (delGCC) and 4PS (delGCCGCC) [15] have evolved independently across multiple isolates (**Fig. 4D**). Taken together, our phylogenomic findings demonstrate that INDELs within *Mtb* SSRs, especially triplet SSRs, are subject to selection, with the distribution of some variants reflecting distinct population- and lineage-specific evolutionary trajectories.

### SSR INDELs are associated with antibiotic resistance in *Mtb* clinical isolates

Next, we utilized a bacterial GWAS approach to identify homoplastic SSR INDELs associated with antibiotic resistance in both *Mtb* populations. We implemented a phylogenetic convergence (phyC) algorithm [9,45], which identifies mutations associated with antibiotic resistance by detecting independent genotype/phenotype co-occurring events across ancestrally reconstructed phylogenetic trees. As expected, DR-associated SNVs were significantly correlated with antibiotic resistance in the two populations (**Fig. 3A-C)**. In addition, we identified 6 SSR INDELs that were significantly associated with isoniazid (INH) resistance in either population (FDR < 0.05) (**Fig. 3A**, *Supplementary Table 3*). Based on the availability of sufficient drug susceptibility data, tests for rifampicin (RIF) and streptomycin (STR) resistance were limited to Peruvian and Vietnamese isolates, respectively, and 4 distinct SSR indels were associated with resistance to these drugs (**Fig. 3B-C**, *Supplementary Table 3*).

**Figure 3.**
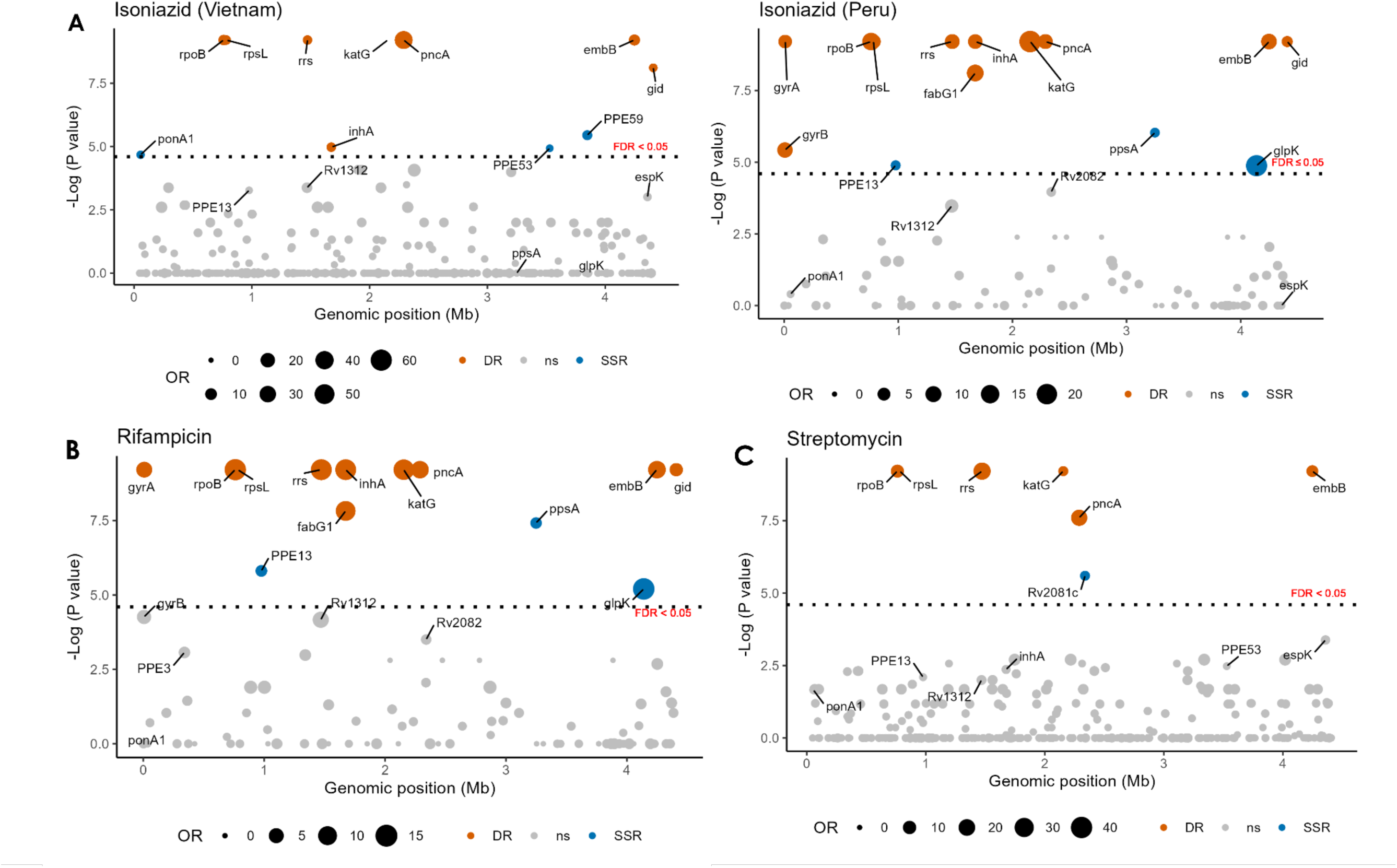
*M. tuberculosis s*imple sequence repeat INDELs are associated with antibiotic resistance. Manhattan plots of the genetic association of repeat indels with (**A**) isoniazid resistance in Vietnamese and Peruvian isolates, **(B**) rifampicin resistance in Peruvian isolates, and (**C**) streptomycin resistance in Vietnamese isolates.

High-confidence associations between INH resistance and SSR INDELs were found in genes previously implicated in antibiotic resistance or tolerance, including *ponA1* and *glpK*. Phase variable *glpK* HT INDELs have previously been linked to multidrug tolerance during TB treatment and are frequently enriched in antibiotic-resistant clinical isolates [12,13,15]. Similarly, SNV mutations in the bifunctional penicillin-binding protein *ponA1* have been associated with RIF resistance, and deletion of this gene reduces RIF potency [9]. The RIF-resistance-associated CCG INDELs that we found are in the triplet SSR that encodes a variable stretch of proline residues within the PonA1-RipA interacting region [24], implicating this interaction in RIF sensitivity.

We also identified resistance associations in genes not previously linked to altered antibiotic susceptibility during TB infection. High-confidence associations with INH resistance in Vietnam were observed for in-frame triplet SSR deletions in both *ppe53* and *ppe59* (**Fig. 3A**). Additionally, frameshifting HT INDELs in *ppe13* and *ppsA* were significantly associated with INH and RIF resistance in Peruvian isolates (**Fig. 3A-B**). Similarly, INDELs in the HT of *Rv2081c* were significantly enriched in STR-resistant isolates from Vietnam (**Fig. 3C**). Consistent with *Mtb* SNVs previously implicated in low-level or intermediate antibiotic resistance [7,8], these resistance-associated SSR INDELs were also identified in drug-susceptible isolates, albeit at significantly lower frequencies (**Fig. 4**). For instance, the estimated 225 independent mutational events that produced *ppe53* in-frame deletions (*ppe53* CGC_del_) resulted in mutations in 34.8% (n=533) of Vietnamese strains, which were significantly enriched in those resistant to INH or STR. Notably, the *ppe53* CGC_del_ variants were virtually absent in L4 strains in both Vietnam and Peru, suggesting strong lineage-specific selection on this INDEL (**Fig. 4**). Overall, we demonstrate that SSR INDELs are significantly associated with antibiotic resistance in a manner that is both drug-specific and population-dependent. While high-level resistance variants are typically absent from susceptible *Mtb* strains, SSR INDELs were present at relatively low frequencies in strains classified as drug sensitive, suggesting a possible role in the evolution of higher-level resistance.

**Figure 4.**
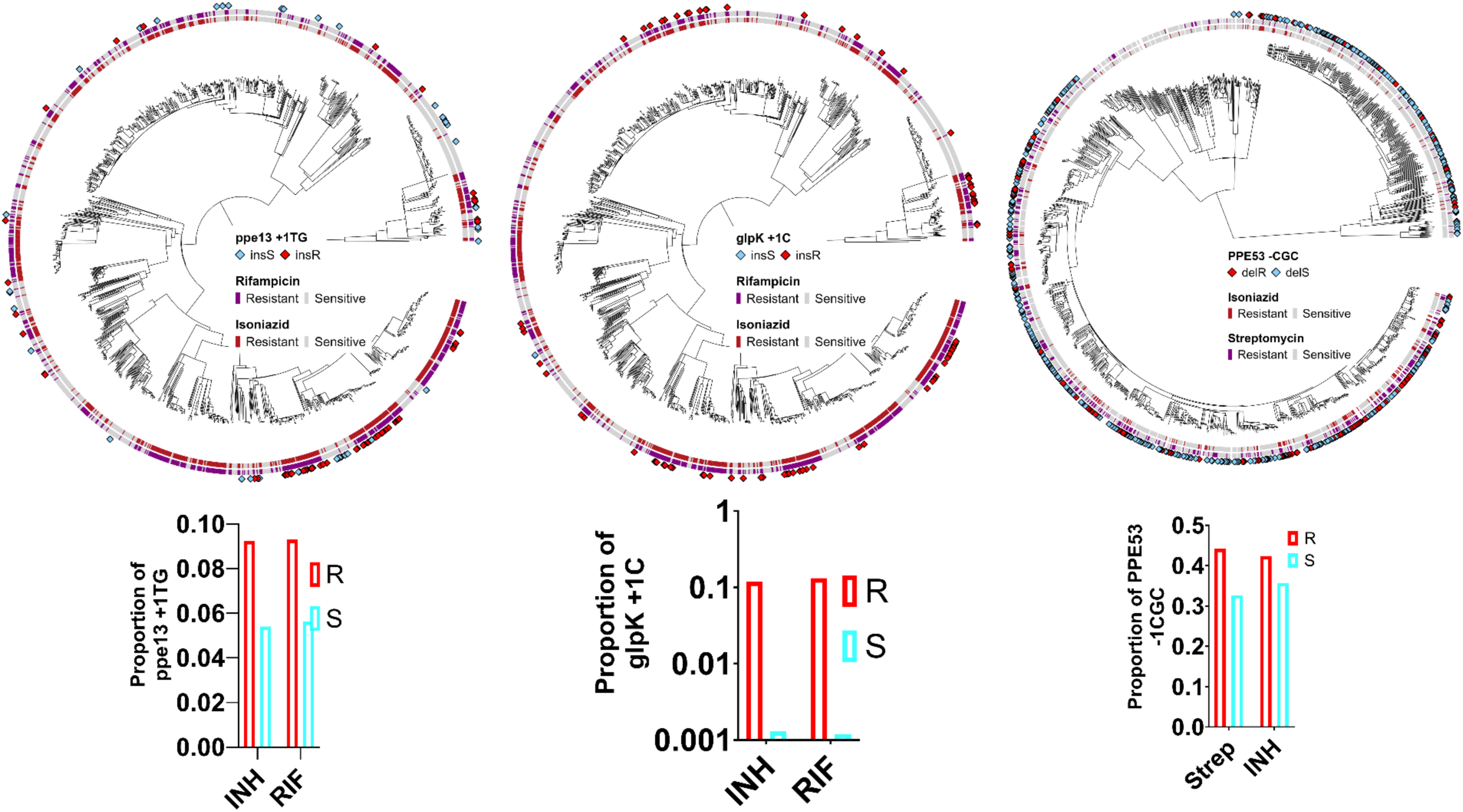
*M. tuberculosis* SSR indels are clinically prevalent and significantly overrepresented in antibiotic-resistant isolates. Phylogenetic trees of the distribution and frequency of *ppe13* and *glpK* HTs insertions and *ppe53* triplet SSR deletion in isoniazid-, rifampicin-, and streptomycin-resistant or sensitive *M. tuberculosis* isolates.

### Engineered SSR variants alter antibiotic susceptibility

To assess the impact of resistance-associated SSR INDELs on drug susceptibility, we engineered a series of these variants and the respective gene knockouts into the H37Rv lab strain background (**Fig. 5A**). Oligo-mediated recombineering was used to introduce clinically prevalent insertions and deletions into the repeat regions of *ppe53* (CGC_del_)*, Rv2081* (-1C)*, Rv2264c* (+1C/+2C)*, espR 5’UTR* (+1G)*, espK* (+1C), and *ponA1* (CCG_ins_). Additionally, oligonucleotide-mediated recombineering followed by the Bxb1 integrase targeting (ORBIT) [46] was used to create gene knockout mutants for all repeat-containing genes except Rv2081c. To enable high-throughput screening of the effects of each SSR or gene knockout mutant on fitness, we introduced a uniquely barcoded, kanamycin selectable integrating plasmid into each strain (**Fig. 5A**). Using this pool of 13 mutants alongside two uniquely barcoded H37Rv wild-type clones, we conducted competition assays to quantify the fitness of each mutant relative to the H37Rv parent. This allowed us to define the relative fitness of each SSR mutant (expressed as normalized mutant abundance) under sub-lethal concentrations (IC10 and IC50) of RIF, INH, and STR.

**Figure 5.**
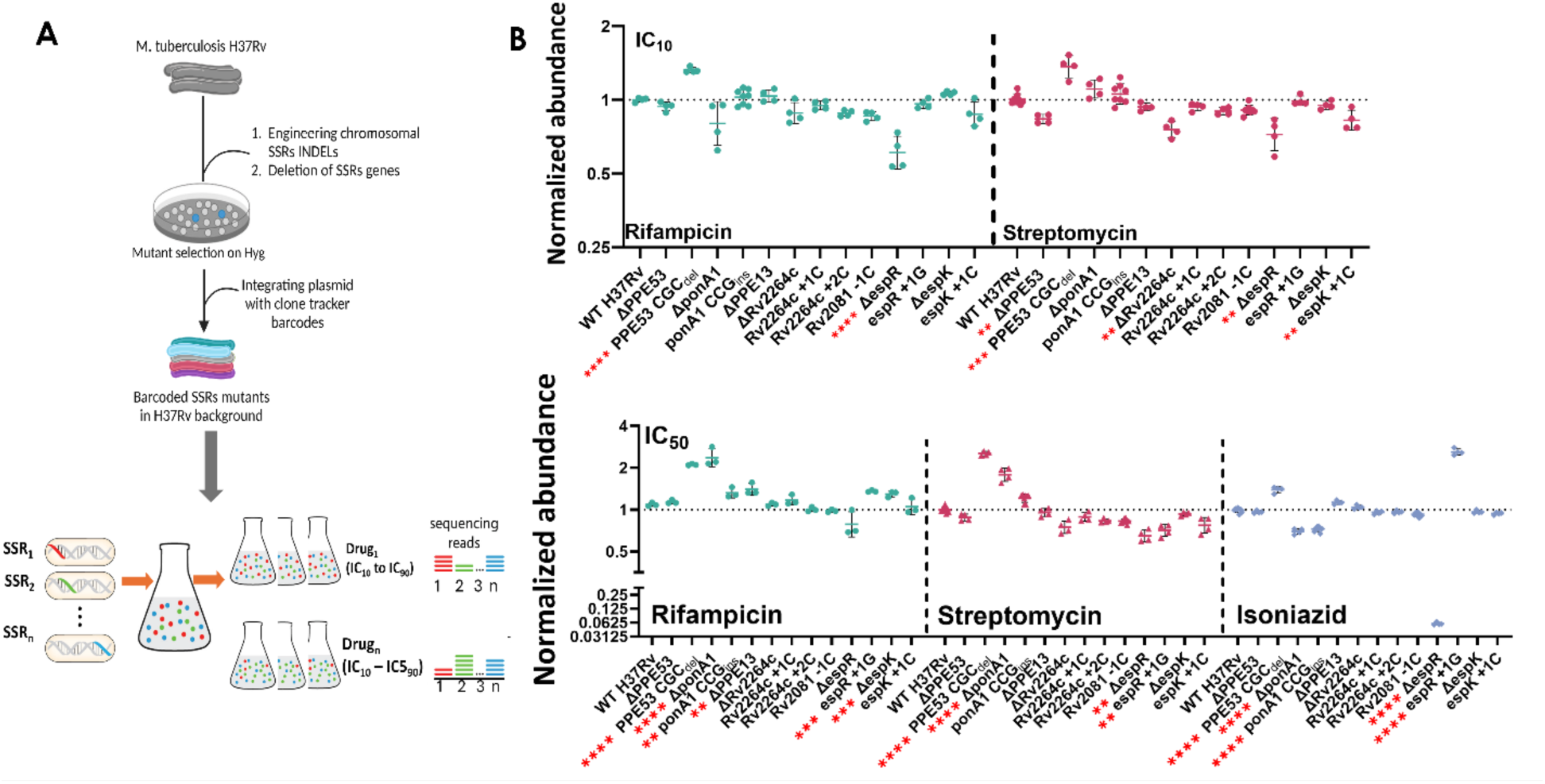
SSR mutants alter *in vitro* antibiotic susceptibility in the H37Rv *M. tuberculosis* background. **(A)** A schematic representation of SSRs mutant generation and barcoding, and the pooled competition assay together with a wild-type H37Rv strain in IC_10_ and IC_50_ of various TB antibiotics. (**B**) Comparison of the normalized abundance of SSR mutants to the barcoded WT H37Rv strain in IC_10_ (top) and IC_50_ (bottom) of rifampicin, isoniazid, and streptomycin. Data shows the mean and the standard error of the mean (SEM) of normalized abundance in biological replicates (n = 4 or 8 if two independent clones are available). Results from Two-way ANOVA with Dunnett’s correction for multiple comparisons; **P < 0.01, ***P < 0.001, and ****P < 0.0001.

We observed variable fitness patterns among the SSR mutants across the antibiotic concentrations (**Fig. 5B**). We previously reported that the *espR* +1G mutation post-transcriptionally upregulates EspR protein levels and reduces INH susceptibility [14]. Consistent with these findings, we found that the *espR* +1G mutant led to a significant decrease in INH susceptibility, while the Δ*espR* mutant displayed increased susceptibility to INH relative to the wild-type strain (**Fig. 5B**). Overall, several of the SSR INDELs with a strong selection signature (**Fig. 2A**), including *Rv2264c, Rv2081c,* and *espK* along with their respective gene knockouts, had no significant impact on antibiotic susceptibility or slightly increased sensitivity to the antibiotics tested in this screen (**Fig. 5B**). However, INDELs in the triplet SSRs of *ponA1* and *ppe53* decreased antibiotic susceptibility across various drugs in the pooled competition assay. Both *ΔponA1* and *ponA1* CCG_ins_ reduced sensitivity to RIF, consistent with a previously characterized knockout mutant [9], leading to a 0.5-to 2.5-fold increase in normalized abundance relative to the wild-type strain. These same mutations increased sensitivity to INH. Most notably, the *ppe53* CGC_del_ mutant, but not the *ppe53* knockout strain, exhibited reduced sensitivity to all three drugs at both doses, consistently increasing *Mtb* fitness across all treatment conditions. (**Fig. 5B**). In summary, these data demonstrate that specific SSR INDELs alter *in vitro* antibiotic susceptibility in *Mtb*. Surprisingly, while in-frame triplet SSR INDELs are expected to have a more modest effect on protein structure than frameshifting INDELs, these variants in PPE53 and PonA1 significantly altered fitness in the presence of antibiotics.

### The *ppe53* CGC_del_ reduces antibiotic susceptibility and disrupts localization of the protein

To further investigate the impact of *ppe53* CGC_del_ on antibiotic susceptibility and address the possibility of unlinked mutations impacting the phenotype of the mutant strain, we repeated the competitive fitness studies using a smaller panel of strains that included a second *ppe53* CGC_del_ mutant independently engineered with the same oligo. Across the three antibiotics, the *ppe53* CGC_del_ variant consistently elevated fitness relative to wild type, whereas deletion of this gene had no effect (**Fig. 6A**). These data suggest that the CGC_del_ mutation does not represent a simple loss of function but rather increases or alters the gene’s effect.

**Figure 6.**
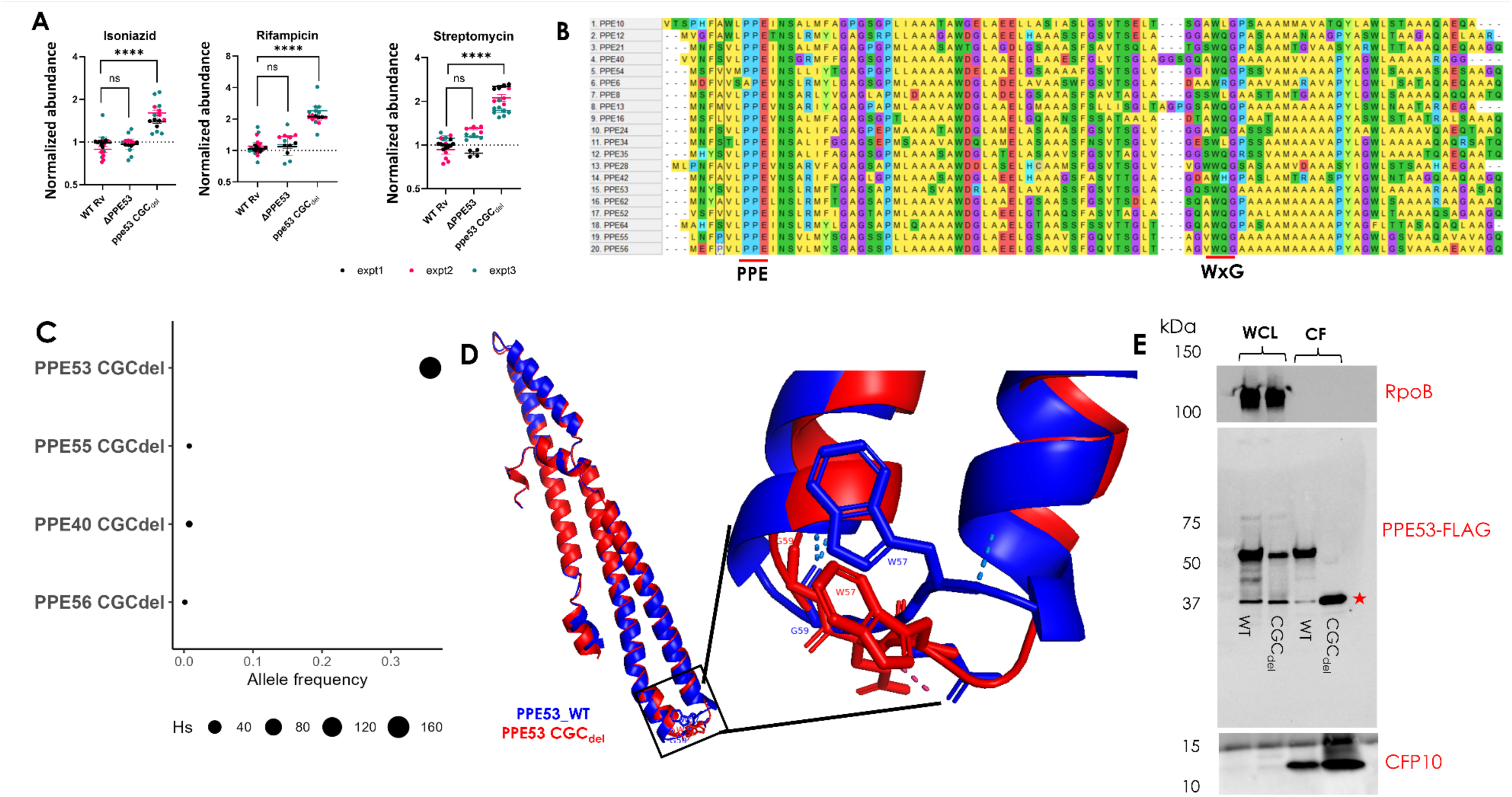
PPE53 triplet deletion alters *in vitro* rifampicin susceptibility and protein’s outer-membrane localization. **(A)** Comparisons of the normalized abundance of the *ppe53* CGC_del_ mutant to WT H37Rv across three independent competition assays in isoniazid, rifampicin, and streptomycin. Experiment 3 included two independent clones of the *ppe53* CGC_del_. Data show the mean and the standard error of the mean (SEM) of normalized abundance in biological replicates (n = 4 or 8 for two independent clones). Results from Two-way ANOVA with Dunnett’s correction for multiple comparisons; **P < 0.01, ***P < 0.001, and ****P < 0.0001. **(B)** Multiple sequence alignment of the N-terminal ends of PPE-MPTR proteins, highlighting the WxG100-adjacent alanine-rich region. **(C)** Frequency of WxG100-adjacent alanine deletions across other PPE-MPTR proteins in 1413 lineage 2 *Mtb* isolates. **(D)** AlphaFold3 model of the N-terminal domains of PPE53 WT and CGC_del_, highlighting the shift in WxG orientation with Ala60 deletion. **(E)** Immunoblots of whole-cell lysates (WCL) and cell filtrate (CF) of PPE53, RpoB, and CFP10 in WT H37Rv and *ppe53* CGC_del_ Mutant.

The *ppe53* triplet repeat encodes an alanine stretch that is conserved within the PPE-Major Polymorphic Tandem Repeat (PPE-MPTR) protein subfamily and lies adjacent to the conserved WxG motif (**Fig. 6B**). This motif is a hallmark of type VII secretion system substrates that serves an important structural role and contributes to the bipartite secretion signal of these proteins [47,48]. The adjacent alanine residues have been termed the “alanine cradle” and are involved in the stabilization of WxG interactions with the type VII secretion machinery [49], suggesting a potential role for this poly-alanine motif in PPE53 outer-membrane localization or secretion. Consistent with this structural model, we found that a WXG-proximal alanine-rich segment is a conserved feature of all members of the PPE-MPTR family. The most similar repeat regions to PPE53 are found in PPE40, PPE55, and PPE56, where the poly-alanine motifs are encoded by triplet repeats identical to PPE53. Despite this similarity, homoplastic INDEL formation in clinical isolates was only observed for PPE53 (**Fig. 6C**), providing further evidence that this SSR in PPE53 is a specific site of selection.

To investigate the biochemical impact of the *ppe53* CGC_del_ mutation, we first used AlphaFold [50] to predict the structures of wild-type and variant proteins. The single alanine deletion reorients the conserved tryptophan (W57) residue away from the helical bundle, disrupting the inter-helical WxG interactions (**Fig. 6D**). Next, to evaluate the effects of this mutation on the outer-membrane localization or secretion of the PPE53 protein, we expressed both the wild-type and CGC_del_ variants of PPE53 in *Mtb*, each fused with an N-terminal FLAG tag. Since the growth of *Mtb* in liquid media supplemented with Tween-80 increases the detection of peripherally cell-associated proteins in the cell culture filtrate [27,51], we reasoned that FLAG immunoblots of whole-cell lysate (WCL) and supernatant (CF) fractions would confirm whether PPE53 is secreted across the plasma membrane and assess the impact of the CGC_del_ mutation on protein localization. We detected two species of the FLAG-tagged wild-type PPE53 in both the WCL and CF, a predominant form corresponding to the 56.6 kDa predicted molecular weight and a second of ∼37 kDa (**Fig. 6E**). As the cytosolic RpoB protein was undetectable in the CF, the presence of wild-type PPE53 in the CF fraction is thus independent of cell lysis and supports our hypothesis that the protein is either secreted or surface-localized. While the PPE53 CGC_del_ gene produced the same two protein products as the wild type in the WCL, their relative abundance differed from the wild type protein, and only the smaller ∼37 kDa product was observed in the CF. This suggests aberrant processing of the secreted mutant protein (**Fig. 6E**). Together, our data show that the in-frame PPE53 deletion decreases drug susceptibility and abrogates the secretion of full-length protein by altering its posttranslational processing. These observations provide a mechanistic basis for the ongoing selection for this variant and its correlation with antibiotic resistance in clinical isolates.

## Discussion

Insertions and deletions within SSRs facilitate transient genetic adaptation to antibiotic and host-induced stress in diverse pathogenic bacteria. In this work, we integrate phylogenomic analyses of clinical isolates with *in vitro* experimentation to identify SSR indels that directly alter drug susceptibility. Using phylogenomic analyses, our work identified SSR INDELs that are highly variable and therefore likely to be under positive selection, those correlated with clinical antibiotic resistance, and those that directly alter drug susceptibility. Most notably, we detected an antibiotic selection signature for in-frame INDELs in triplet SSRs across cell wall and membrane-associated *Mtb* genes, and that a clinically prevalent in-frame deletion in *ppe53* confers intermediate antibiotic resistance by disrupting the processing and secretion of this protein. Our results add to previous studies that identified *Mtb* mutations linked to intermediate antibiotic resistance or tolerance and highlight the significance of non-frameshifting triplet SSRs in *Mtb* adaptation to antibiotic stress.

While previous GWAS of *Mtb* SSR INDELs focused on global collections of isolates to map the selection signatures [15], we narrowed our analysis to two distinct populations to resolve nuanced inter-population differences in the frequency and selection of these INDELs. For several SSR INDELS, substantial differences were observed between populations in both the degree of homoplasy and allele frequency. For example, the frequency of the *glpK* HT insertions, previously linked to *Mtb* antibiotic recalcitrance, was significantly higher in Peruvian relative to Vietnamese strains. This could be explained by variation in pre-sequencing sample culturing techniques, particularly the routine use of glycerol-based culture media, which selects against *glpK* phase variants [13,52]. For other variants, these differences may stem from differences in TB control programs, such as a history of streptomycin use, higher incidence of rifampicin- or multidrug-resistant TB in Peru [53], or the unique bacterial genetic backgrounds.

Overall, we observed a strong selection signature on HT INDELs across multiple *Mtb* genes, including those previously shown to be enriched in antibiotic-resistant *Mtb* strains or to alter *in vitro* drug susceptibility [12,14,15]. Additional frameshifting HT variants with high homoplasy scores included those in *ppe13, bioF2, espK,* and *Rv2081c*, all of which have previously been shown to undergo selection for SNV accumulation in *Mtb* isolates [15]. Interestingly, several of the most homoplastic SSR INDELs did not alter *in vitro* antibiotic sensitivity. For instance, insertions within the *espK* HT locus, a region exhibiting a strong selection signature in this and prior work [15], either increased bacterial susceptibility or had no detectable impact on antibiotic efficacy. EspK serves as a chaperone for ESX-1-mediated secretion of various substrates, including EspB [54], and is frequently mutated during *Mtb* infection [55], highlighting a potential role in *Mtb* virulence instead of antibiotic adaptation. Similarly, indels within both *Rv2264c* and *Rv2081c* HTs were among the most homoplastic SSR variants, and previous studies have shown a robust correlation between *Rv2264c* HT mutants and clinical resistance to multiple antibiotics [15]. However, the three *Rv2264c* mutants and the single *Rv2081c* mutant evaluated in our pooled competition assays failed to provide a significant fitness advantage in the presence of the tested drugs (**Fig. 5**). Rv2264c is predicted to be an outer-membrane or cell wall-associated protein containing potential B-cell immunogenic epitopes [56,57], while Rv2081c ranks among the most abundant *Mtb* mRNA transcripts across diverse culture conditions [58]. The lack of significant change in antibiotic susceptibility in our assays suggests that the selection of these variants may be driven by pressures other than antibiotics, that they exert drug-related effects not captured by our simple *in vitro* assays, or that their effects depend on the strain’s genetic background.

Among the most homoplastic SSR loci were several within PPE proteins, an enigmatic paralog family expanded in pathogenic mycobacteria [59]. Notably, some of the variations in these PPE genes correlated with clinical drug resistance and caused *in vitro* intermediate resistance. INDELs in the contiguous 9G/8T HT of *ppe13* were among the most homoplastic and significantly correlated with INH and RIF resistance in Peruvian strains. Consistent with this observation, past genomic analyses have linked *ppe13* SNVs with antibiotic resistance, and also inferred strong selective pressure on the gene based on high nonsynonymous to synonymous polymorphism ratios (pNpS) in drug-resistant cohorts [29,31]. Our results also identified triplet SSRs in other PPE genes, including *ppe26, ppe60, ppe53,* and *ppe64,* displaying a pattern of ongoing selection. Recent studies suggest that PE/PPE heterodimers form hydrophilic pores or channels that mediate nutrient transport and molecular permeability across the mycobacterial outer membrane (MOM) [28,30,60]. PPE13, PPE53, and PPE64 were computationally predicted to be MOM-associated [56], confirming their exposure at the host-pathogen interface and highlighting their role in antibiotic- or host-driven selection. Indeed, prior studies have implicated PPE13 in clinical antibiotic resistance [29], PPE64 in nutrient uptake [61], and PPE60 in within-host adaptation [62], underscoring the potential role of PPE-associated SSRs in mediating mycobacterial responses to a changing host environment.

The selection pattern of in-frame INDELs in the triplet SSRs, particularly those within genes associated with cell wall biogenesis or PPEs predicted to be outer membrane proteins, emerged as the most striking finding in our study. Triplet SSRs translate into amino acid homopeptide stretches, which can contribute to protein structure, protein-protein interactions, and surface localization [22,23]. While expansions and contractions of these genomic features have been implicated in multiple human disorders [63], their functional consequences in bacterial adaptation to stress are less clear. In the peptidoglycan biosynthetic protein PonA1, expansions and contractions in a proline stretch of the C-terminal domain previously implicated in PonA1-RipA interactions [24], were homoplastic and associated with antibiotic resistance in our study. Variability in the length of this proline stretch has also been linked to alterations in mycobacterial growth rate and cell length [25]. Notably, we found that the frequency and selection patterns in the *ponA1* triplet SSR varied substantially across *Mtb* lineages, a pattern driven primarily by differences in the ancestral polyproline lengths across *Mtb* lineages. While beyond the scope of this work, recent advances in cytological profiling [64] could be leveraged to link variations in this polyproline stretch with morphological differences across *Mtb* lineages.

The most prominent in-frame triplet SSR variant in our study was the PPE53 CGC_del_, which exhibited a strong selection signature, correlated with clinical resistance, and decreased *in vitro* susceptibility to multiple antibiotics. Despite exhibiting a low pNpS ratio for SNVs, characteristic of purifying selection [31], PPE53 has been linked to adaptation in lipid-rich environments [65], and transposon mutants in this gene display β-lactam resistance [66]. In *M. marinum,* PPE53 was identified as a virulence factor required for infection of and growth in macrophages [67,68]. Even though the clinically prevalent *ppe53* CGC_del_ variant exhibited intermediate antibiotic resistance in the H37Rv strain, the knockout mutant showed no effect on antibiotic susceptibility. We hypothesize that the CGC_del_ mutation, rather than causing a loss of function, results in a modification of the PPE53 protein activity.

Biochemical analysis of the PPE53 protein revealed two fragments, corresponding to the full-length protein (56.6 kDa) and a smaller fragment (∼37 kDa), in both the WCL and CF of the wild type, whereas only the smaller fragment was detected in the CF of the CGC_del_ mutant. As the majority of PPE-MPTRs, including PPE53, are predicted to be substrates of the ESX-5 type VII secretion system (T7SS) [69–71], we reason that deletion in this polyalanine stretch potentially disrupts key interactions between PPE53 and components of T7SS machinery, impairing PPE53 recognition, processing, and translocation to the MOM. This interpretation is supported by the observed reorientation and loss of key interactions by W57 and G59 residues, both part of the WxG domain, in the *ppe53* CGC_del_ mutant relative to the wild-type protein. The recognition, processing, and translocation of ESX-5 substrates is mediated by conserved bipartite secretion signals, including the N-terminal WxG domain [72,73]. The *ppe53* CGC_del_ mutation maps to an alanine-rich region adjacent to the WxG domain, which is conserved across other PPE-MPTR proteins. Previous studies have implicated this ‘alanine cradle’ in the stabilization of key interactions between the ESX-1 T7SS complex and its substrates, enabling the proper recognition and export of proteins across the mycomembrane [49]. Thus, our observation of defective surface localization of the PPE53 protein in the CGC_del_ mutant could be explained by disrupted WxG interactions in this mutant, highlighting an unrecognized role for alanine homopeptide variation in the structural organization, recognition, processing, and secretion of PE/PPE substrates.

Overall, our study mapped selection signatures across *Mtb* SSR INDELs, uncovering a role for triplet SSR variation in intermediate antibiotic resistance. More specifically, we identified an in-frame INDEL in *ppe53* that broadly reduces the potency of multiple drug classes. While these changes in drug efficacy are modest, even small MIC increases to RIF can increase the rate of treatment failure [10]. The high prevalence of this mutation in L2 strains, including those that are classified as drug-susceptible, suggests that its presence may commonly promote the evolution of higher-level resistance. Our work highlights the significance of triplet repeats in *Mtb*’s adaptation to drug stress during infection, warranting further genomic and experimental validation to more fully elucidate their roles in the evolutionary path to antibiotic resistance.

## Materials and Methods

### Sequence data processing, mapping, and variant calling

We curated a list of publicly available and unpublished *Mtb* whole-genome sequencing data from Vietnamese and Peruvian isolates [32,33]. The list of SRA project numbers, accession IDs, drug susceptibility profiles, and demographic information is summarized in *Supplementary Table 1*. The SRAtoolkit (v3.0.0) *prefetch* and *fastq-dump* functions were used to download *sra* and paired-end *fastq* files while maintaining the original file format. All Sequence reads were first processed with *Sickle* (v1.33) to trim low-quality 3’ and 5 ′ end reads while retaining reads with length > 100 and Phred score > 30 for subsequent analyses (*sickle pe -t sanger -q 30 -l 100*). We used the inferred *MTB*C ancestral genome of the most recent common ancestor (MRCA) [74] as a reference, and all trimmed reads were mapped using Bowtie2 (v2.5.0) [75]. Samtools (v1.16.1) was used to convert the resulting sequence alignment map (SAM) files to binary alignment map (BAM) files, followed by processing steps to fill in mate coordinates (*samtools fixmate*), sort (*samtools sort*), and mark duplicate reads (*samtools markdup*), compute sequence depth and breadth (*samtools depth*), and index processed BAM files (*samtools index*). To enhance the accuracy of indel calls in repetitive regions, we only retained samples with an average sequencing depth above 30X and a mapping rate above 95% for downstream analyses.

To identify the SSR coordinates in the reference genome, we used the *TRFinder* (v4.09.1) algorithm with default parameters [76]. We then implemented an indel genotyping (*gatk HaplotypeCaller –R –I [BAM.list] –ploidy 1 -L SSRs.bed –o [vcf]*) and re-genotyping (*gatk HaplotypeCaller -I [BAM.list] -R --alleles [vcf] -gt_mode GENOTYPE_GIVEN_ALLELES -ploidy 1 -o [regenovcf]* algorithms using GATK v4.3.0 to call all high-quality insertions and deletions mapping to the specified repeat coordinates. We applied hard filtering on all indels as previously described [36], retaining high-quality short indels (<= 10 base deletion or insertion) with minimum coverage > 25 reads at the position, mean base quality > 25, and a mean mapping quality > 30. All SNVs aligning to coding and intergenic regions of genes previously associated with antibiotic resistance were also called. VCF files were processed using BCFtools (v0.1.16), excluding samples with no variant calls in more than 25% of SSR positions and positions with missing variant calls in over 25% of individuals, and subsequently annotated using SnpEff [77] using *the Mycobacterium tuberculosis H37Rv* database as the reference.

### Phylogenetic Analysis and Homoplasy Score Estimation

We constructed the core SNV-based phylogenetic trees for strains from the two populations as previously described [6]. Briefly, all fixed SNPs were genotyped independently for Peruvian and Vietnamese isolates using *VarScan* (v2.3.9) with the strand bias on, retaining SNVs with a frequency of > 90% and supported by at least 10 reads. SNPs mapping to repetitive regions of the MRCA genome, including PE/PPE genes, prophages, and mobile genetic elements, were excluded. All SNP locations were subsequently merged into a non-redundant consensus list and recalled using the *VarScan* (v2.3.9) *mpileup2cns* function. This process generates a FASTA file containing all core SNVs across isolates while excluding monomorphic loci in isolates and the MRCA reference. Nucleotide positions with no calls in more than 5% of isolates were excluded, and all FASTA files were subsequently merged to produce a gap-free multi-sequence alignment file. We then used the alignments with the remaining polymorphic positions from all strains in the two populations to construct the trees using RAxML-ng (v1.2.1) [78], with the GTRgamma substitution model and at least 200 bootstrapping replicates. We included a strain of *Mycobacterium canettii*, an extant progenitor of the *MTB*C [79], as an outgroup for both trees. All phylogenies were annotated and visualized using *ggtree* (v3.8.2).

To quantify the number of independent variant occurrences (homoplasy scores, Hs), we used TopDIS, a recently developed algorithm that integrates variant VCF files, phylogenetic trees, and isolate metadata to infer convergent evolution [15]. We first constructed a *position_variant* and *isolate_ID* matrix independently for the two populations, decoding positions with missing calls as *99.* The space length setting was set at 3 [15] for both SSR indels and DR-associated SNPs in the two populations. All variants with Hs < 1 were excluded from downstream analysis.

### Tests for antibiotic-resistance associations

To identify repeat indels significantly associated with antibiotic resistance during TB infection, we used a recently published tool (Hogwash v1.2.6) [45] that implements a phylogenetic convergence-based bacterial GWAS protocol [9]. With each phylogenetic tree and trait (antibiotic resistance), a binary ancestral reconstruction is done for each phenotype and genotype. This excludes all tree edges with low bootstrap support (<95%), long tree lengths, or a maximum likelihood < 0.875 (low ancestral reconstruction support) [45]. Hogwash then implements the phyC algorithm, identifying the overlap of the phenotype (antibiotic resistance) with the genotype transition on the phylogenetic tree. To enhance the power of the association tests, we utilized the post-ancestral reconstruction grouping feature of the algorithm, clustering all variants from the same gene before identifying those significantly associated with antibiotic resistance. The algorithm generates a null distribution and an empirical *P-*value comparison of the observed vs null phenotype-genotype transitions. To enhance the sensitivity of the genomic association test, we used pheno_present/pheno_absent and geno_transition/geno_not_transition contingency tables to estimate the effect size (odds ratio) for each association. Known DR SNVs were included in the association tests to serve as controls in the analyses.

### Growth conditions, strains, and plasmid construction

The standard Middlebrook 7H9 media (7H9 salts, 0.2% glycerol, 0.05 Tween-80, and 10% OADC) was used in all cultures unless otherwise indicated. A modified 7H9 broth medium, with 0.05% tyloxapol and 0.1mM propionate, was used for the outgrowth of colonies to ensure the selection of phthiocerol dimycocerosate (PDIM) positive cells [80]. For protein extraction, *Mtb* cells were grown in the salt-based minimal media (asparagine (0.5 g/l), KH_2_PO_4_ (1.0 g/l), Na_2_HPO_4_ (2.5 g/l), ferric chloride (50 mg/l), MgSO_4_ ·7H_2_0 (0.5 g/l), CaCl_2_ (0.5 g/l), ZnSO_4_ (0.1 mg/L)) with 0.2% glycerol and 0.05% Tween-80. For the outgrowth of *Mtb* cells on solid media, 7H10 agar supplemented with 10% OADC and 0.5% glycerol (in the presence of the respective antibiotics) was used. All experimental procedures involving the virulent *Mtb* H37Rv strains and all other derivative mutants were conducted in adherence to the CDC-NIH biosafety guidelines in biosafety level 3 (BSL-3) laboratories.

All strains, including the chromosomal SSR indels or gene knockouts, were done in the *Mtb* H37Rv background using oligo-mediated recombineering and ORBIT [46] and adhered to existing NIH Guidelines for research with recombinant DNA. Briefly, to introduce repeat insertions and deletions in the respective gene repeats, we designed 70-base-long oligonucleotides containing the insertion or deletion of interest and sequences that overlap the region (Supplementary Table 4). H37Rv strain with integrated plasmids expressing the kanamycin-selectable phage Che9c RecT from the inducible P_tet_ promoter (pKM402) and the L5 integrating vector with a defective hygromycin (early termination codon) (pKM 427) was inoculated in standard 7H9 broth, with anhydrotetracycline (ATc) induction of RecT expression. The SSR indel-specific and the 70-base Hyg repair oligo were co-electroporated in electrocompetent H37Rv cells and selected on hygromycin (50 µg/ml) 7H10 plates. For each transformation, colonies were screened for respective SSR indels by Sanger sequencing, followed by re-streaking and curing of the pKM402 plasmid.

To generate respective gene knockouts, we created 188bp ultra-mers, containing the 48-base *attP* site flanked by 70 bases specific to the gene on both sides (Supplementary Table 4). H37Rv *Mtb* cells expressing the RecT and Bxb1 integrase under the control of the Ptet promoter (pKM461) were grown in standard 7H9 media, with ATc induction 24 hours before the transformation. 1µg of the ORBIT oligo was co-electroporated with 200ng of the non-replicating payload plasmid (pKM464), overgrown overnight, and selected in 7H10 plates with 50µg/ml of hygromycin. Successful colonies were screened by PCR and Sanger sequencing of the gene/hyg-out and gene/ori-out junctions as previously described [46].

To uniquely barcode all the chromosomal repeat mutants and knockout strains, we first cured all the strains of the kanamycin-resistant plasmid (pKM402 and pKM461) followed by electroporation with a Tweety-integrating, kanamycin-selectable plasmid (pKP1341) with a unique 20-mer barcode. Successful transformants were selected in 7H10 solid agar with 25µg/ml kanamycin, and outgrown in modified 7H9 with 0.05% tyloxapol, 0.1mM propionate, and 0.2% glycerol. All barcoded strains were validated by Sanger sequencing of both the barcode junction on the plasmid and the respective SSR mutations or gene deletions.

To express the FLAG-tagged PPE53 (wild-type and CGC_del_) genes, we used anchored primers (Supplementary Table 4) to amplify the *ppe53* gene from genomic DNA extracted from *Mtb* H37Rv wild-type and mutant strains by PCR. This was followed by the two-step Gateway cloning of the amplicons, creating two independent plasmid constructs (WT and CGC_del_), each containing an N-terminal 3X-FLAG tag driven by a synthetic promoter in the pDE43-MCK/MCZ vector. All plasmid constructs were confirmed by Sanger sequencing.

### Secretion analysis by immunoblotting

The secretion analysis of PPE53 (WT and CGC_del_) was done as described earlier, with a few modifications [81]. Briefly, both strains were first grown to the mid-logarithmic phase in standard Middlebrook 7H9 broth in the presence of appropriate antibiotics. Cells were subsequently washed and resuspended in the salt-based minimal media supplemented with 0.2% glycerol and 0.05% Tween-80 at an OD_600_ of 0.35 and incubated for 48 hours. Cells were harvested by centrifugation, and the pellet was washed in phosphate-buffered saline (PBS) and resuspended in the mycobacterial protein extraction buffer (1M Tris-Cl (pH 7.5), 0.5M EDTA, 20% SDS, and Roche protease inhibitor cocktail tablet in sterile water), followed by bead beating and centrifugation. The supernatant was filtered twice through the 0.22µm filter and concentrated in the Amicon Ultra-3K conical tubes (Merck Millipore, UFC900324) by centrifugation. We measure protein concentrations using the Pierce BCA Protein Assay kit (Thermo Scientific, 23225) with necessary dilutions in the protein extraction buffer. SDS-PAGE western blots were stained with HRP2-conjugated anti-FLAG monoclonal antibody (Millipore Sigma, A8592), and rabbit polyclonal antibodies against RpoB (BioLegend, 663905) and ESAT6 (Sigma, SAB4701024).

### *In vitro* competition experiments

For competition assays, all barcoded repeat strains were individually inoculated in 7H9 media and grown to mid-log phase (OD_600_ ∼ 0.6). All strains were then pooled together at equivalent OD_600_ and then inoculated in quadruplets in fresh 10ml liquid media at OD_600_ ∼ 0.005 containing antibiotics at indicated concentrations or no antibiotics for growth controls. All cultures were incubated at 37 degrees with constant shaking for 6 days, followed by OD_600_ measurements. To quantify the relative fitness of individual repeat mutants in the pool in different antibiotics, genomic DNA (gDNA) was extracted as previously described [8]. Briefly, cells were spun down, resuspended in 1ml of TE buffer, and added into bead-beating tubes with 500µl of 25:24:1 phenol:chloroform:isoamyl alcohol solution and 300µl equivalent 0.1mm Zirconia beads. This was followed by bead-beating four times for 45 seconds at maximum speed, followed by centrifugation at 15,000 rpm for 10 minutes. The aqueous layer was then transferred to new tubes containing an equal volume of 1:1 phenol: chloroform before being safely transferred out of the BSL3 facility. DNA was cleaned using the phenol: chloroform method as previously described [82], precipitated in 1/10 volume 3M sodium acetate and 1 volume isopropanol, and the concentration was determined using Nanodrop 2000/2000c.

For Illumina sequencing library preparation, the plasmid barcode junction was amplified from 40 µg gDNA with 22 cycles using the Q5 High Fidelity 2X master mix (NEB, M0492L) in a reaction containing 4µl of gDNA, 6µl of 10µm primer pairs, 20µl of Q5 High Fidelity 2X master mix, and 4µl of DNase-free molecular grade water. The primers were designed to include TruSeq dual index, and a custom stagger sequence was included in the reverse primer to improve the base diversity for Illumina sequencing. Libraries were individually cleaned using the AmpureXP beads, followed by library quantification using the Qubit 1x dsDNA HS assay kit (Invitrogen, Q33231). All libraries were pooled together at equimolar concentration, and the concentration was confirmed using the Kappa Library Quantification Kit (Roche, KK4824). The final sequencing library was spiked with 5-10% of phiX spike-in (NextSeq phiX control Kit, Illumina) to improve the sequencing diversity. This was followed by Illumina deep sequencing on the Illumina NextSeq 1000/2000 platform (single-end, dual-indexed, 1x100 cycles).

Barcode counts were extracted from raw sequencing files using a custom Bash script, ensuring no mismatches in the barcode or intervening 10bp sequences. To estimate the barcode abundance in biological replicates across experiments, the raw read counts were first expressed as a fraction of the total library size and then normalized by dividing each strain’s fraction by the average fraction of WT strains, setting WT replicates to 1. To further account for differences in the input library (day 0), the normalized strain abundance was divided by the input library’s normalized barcode abundance. Strain abundance data for biological replicates are plotted as the mean and standard deviation.

## Acknowledgements

We appreciate members of the Sassetti Lab at UMass Chan Medical School for the insightful discussions that shaped this project. We also acknowledge the lab of Megan Murray at Harvard Medical School, especially Chuan-Chin Huang, for help with the Peruvian WGS dataset. This work was supported by NIH/NIAID grants AI62598 and AI143575 to CMS.

## Conflicts of interest

Authors declare no conflicts of interest

## Data availability

Whole genome sequencing data analyzed in this manuscript were sourced from previously published studies and are publicly available. Additional Accession IDs and BioProject details are provided in supplemental details.

## Supporting Information

**Figure S1.**
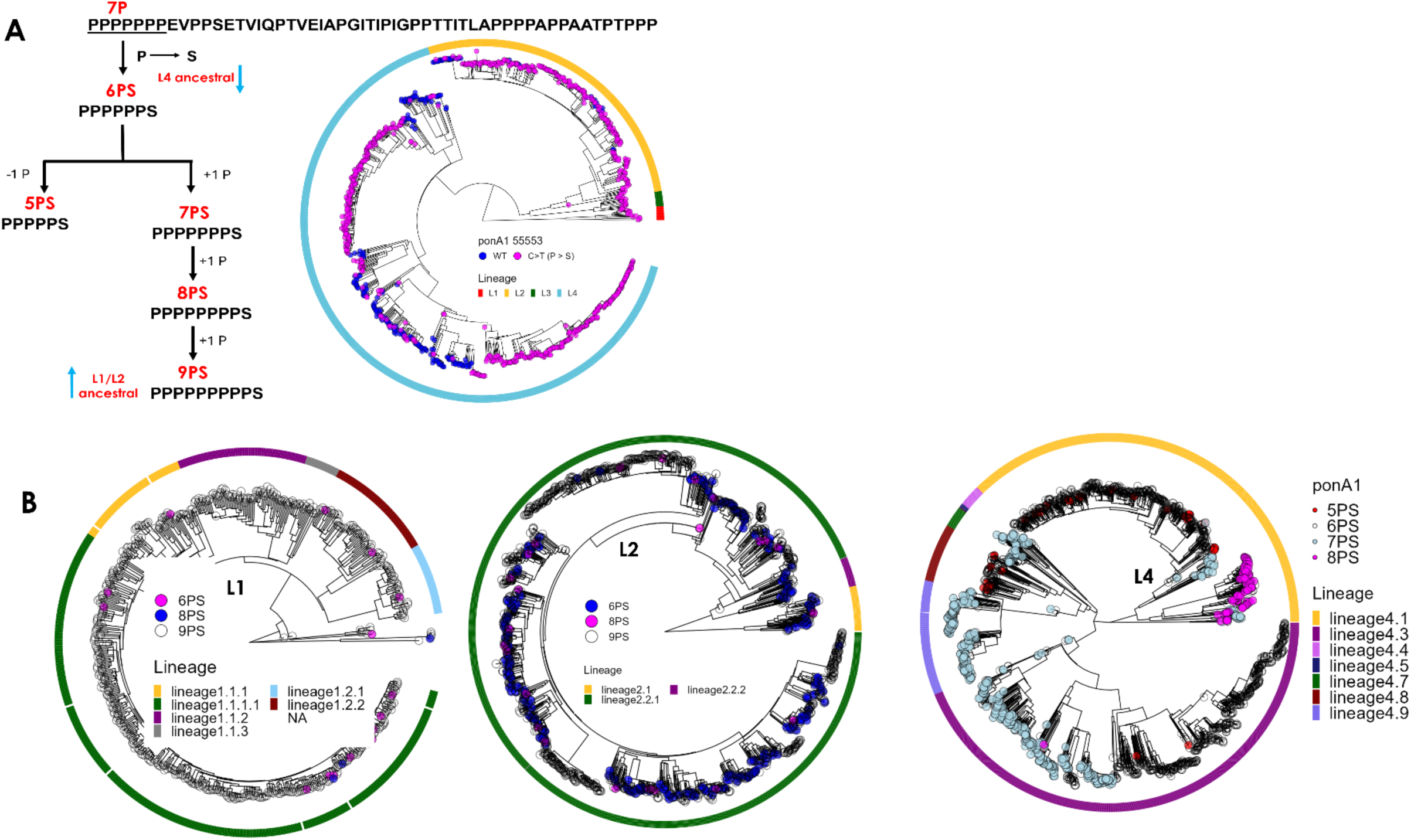
Distribution of *ponA1* proline repeats in L1, L2, and L4 of *M. tuberculosis.* **(A)** Schema of *ponA1* proline repeats (left) and phylogenetic distribution of the fixed CCG > TCG (Pro > Ser) across 4 *Mtb* lineages. **(B)** Phylogenetic relationship of proline repeats across L1, L2, and L4 of *Mtb*.

**Table S1:** Information on publicly available WGS of Mtb isolates used in this work

**Table S2:** Homoplasy scores of SSR variants and DR SNVs in Mtb isolates from Peru and Vietnam

**Table S3:** Data on the antibiotic resistance association test on Mtb isolates from Vietnam and Peru.

**Table S4:** Oligos and primers used in this study

## Notes

### Competing Interest Statement

The authors have declared no competing interest.

## References

1. Global Tuberculosis Report 2025. [cited 30 Apr 2026]. Available: https://www.who.int/teams/global-programme-on-tuberculosis-and-lung-health/tb-reports/global-tuberculosis-report-2025

2. Catalogue of mutations in Mycobacterium tuberculosis complex and their association with drug resistance, 2nd ed. World Health Organization; 15 Nov 2023 [cited 30 Apr 2026]. Available: https://www.who.int/publications/i/item/9789240082410

3. Walker TM, Miotto P, Köser CU, Fowler PW, Knaggs J, Iqbal Z, et al. The 2021 WHO catalogue of Mycobacterium tuberculosis complex mutations associated with drug resistance: A genotypic analysis. Lancet Microbe. 2022;3: e265–e273.

4. García-Marín AM, Cancino-Muñoz I, Torres-Puente M, Villamayor LM, Borrás R, Borrás-Máñez M, et al. Role of the first WHO mutation catalogue in the diagnosis of antibiotic resistance in Mycobacterium tuberculosis in the Valencia Region, Spain: a retrospective genomic analysis. Lancet Microbe. 2024;5: e43–e51.

5. Worakitchanon W, Yanai H, Piboonsiri P, Miyahara R, Nedsuwan S, Imsanguan W, et al. Comprehensive analysis of Mycobacterium tuberculosis genomes reveals genetic variations in bacterial virulence. Cell Host Microbe. 2024;32: 1972–1987.e6.

6. Liu Q, Zhu J, Dulberger CL, Stanley S, Wilson S, Chung ES, et al. Tuberculosis treatment failure associated with evolution of antibiotic resilience. Science. 2022;378: 1111–1118.

7. Hicks ND, Yang J, Zhang X, Zhao B, Grad YH, Liu L, et al. Clinically prevalent mutations in Mycobacterium tuberculosis alter propionate metabolism and mediate multidrug tolerance. Nat Microbiol. 2018;3: 1032–1042.

8. Hicks ND, Giffen SR, Culviner PH, Chao MC, Dulberger CL, Liu Q, et al. Mutations in dnaA and a cryptic interaction site increase drug resistance in Mycobacterium tuberculosis. PLoS Pathog. 2020;16: e1009063.

9. Farhat MR, Shapiro BJ, Kieser KJ, Sultana R, Jacobson KR, Victor TC, et al. Genomic analysis identifies targets of convergent positive selection in drug-resistant Mycobacterium tuberculosis. Nat Genet. 2013;45: 1183–1189.

10. Colangeli R, Jedrey H, Kim S, Connell R, Ma S, Chippada Venkata UD, et al. Bacterial Factors That Predict Relapse after Tuberculosis Therapy. N Engl J Med. 2018;379: 823–833.

11. Meehan CJ, Goig GA, Kohl TA, Verboven L, Dippenaar A, Ezewudo M, et al. Whole genome sequencing of Mycobacterium tuberculosis: current standards and open issues. Nat Rev Microbiol. 2019;17: 533–545.

12. Bellerose MM, Baek S-H, Huang C-C, Moss CE, Koh E-I, Proulx MK, et al. Common Variants in the Glycerol Kinase Gene Reduce Tuberculosis Drug Efficacy. MBio. 2019;10. doi:10.1128/mBio.00663-19

13. Safi H, Gopal P, Lingaraju S, Ma S, Levine C, Dartois V, et al. Phase variation in Mycobacterium tuberculosis glpK produces transiently heritable drug tolerance. Proc Natl Acad Sci U S A. 2019;116: 19665–19674.

14. Luna MJ, Oluoch PO, Miao J, Culviner P, Papavinasasundaram K, Jaecklein E, et al. Frequently arising ESX-1-associated phase variants influence Mycobacterium tuberculosis fitness in the presence of host and antibiotic pressures. MBio. 2025; e0376224.

15. Vargas R Jr, Luna MJ, Freschi L, Marin M, Froom R, Murphy KC, et al. Phase variation as a major mechanism of adaptation in complex. Proc Natl Acad Sci U S A. 2023;120: e2301394120.

16. Moxon R, Bayliss C, Hood D. Bacterial contingency loci: the role of simple sequence DNA repeats in bacterial adaptation. Annu Rev Genet. 2006;40: 307– 333.

17. Ellegren H. Microsatellites: simple sequences with complex evolution. Nat Rev Genet. 2004;5: 435–445.

18. Rodríguez-Pastor R, Knossow N, Shahar N, Hasik AZ, Deatherage DE, Gutiérrez R, et al. Pathogen contingency loci and the evolution of host specificity: Simple sequence repeats mediate Bartonella adaptation to a wild rodent host. PLoS Pathog. 2024;20: e1012591.

19. Jerome JP, Bell JA, Plovanich-Jones AE, Barrick JE, Brown CT, Mansfield LS. Standing genetic variation in contingency loci drives the rapid adaptation of Campylobacter jejuni to a novel host. PLoS One. 2011;6: e16399.

20. Yu Q, Mortimer TD, Blomqvist SOP, Bowcutt B, Helekal D, Palace SG, et al. Diversity and evolution of a phase-variable multi-locus antigen in Neisseria gonorrhoeae. PLoS Pathog. 2026;22: e1013962.

21. Sreenu VB, Kumar P, Nagaraju J, Nagarajam HA. Simple sequence repeats in mycobacterial genomes. J Biosci. 2007;32: 3–15.

22. Chavali S, Singh AK, Santhanam B, Babu MM. Amino acid homorepeats in proteins. Nat Rev Chem. 2020;4: 420–434.

23. Faux NG, Bottomley SP, Lesk AM, Irving JA, Morrison JR, de la Banda MG, et al. Functional insights from the distribution and role of homopeptide repeat-containing proteins. Genome Res. 2005;15: 537–551.

24. Hett EC, Chao MC, Rubin EJ. Interaction and modulation of two antagonistic cell wall enzymes of mycobacteria. PLoS Pathog. 2010;6: e1001020.

25. Gao B, Wang J, Huang J, Huang X, Sha W, Qin L. The dynamic region of the peptidoglycan synthase gene, Rv0050, induces the growth rate and morphologic heterogeneity in Mycobacteria. Infect Genet Evol. 2019;72: 86–92.

26. Otsuka Y, Parniewski P, Zwolska Z, Kai M, Fujino T, Kirikae F, et al. Characterization of a trinucleotide repeat sequence (CGG)5 and potential use in restriction fragment length polymorphism typing of Mycobacterium tuberculosis. J Clin Microbiol. 2004;42: 3538–3548.

27. Ates LS, van der Woude AD, Bestebroer J, van Stempvoort G, Musters RJP, Garcia-Vallejo JJ, et al. The ESX-5 System of Pathogenic Mycobacteria Is Involved In Capsule Integrity and Virulence through Its Substrate PPE10. PLoS Pathog. 2016;12: e1005696.

28. Boradia V, Frando A, Grundner C. The Mycobacterium tuberculosis PE15/PPE20 complex transports calcium across the outer membrane. PLoS Biol. 2022;20: e3001906.

29. Boradia V, Chen J, Frando A, Clark LV, Grundner C. PE/PPE proteins contribute to Mycobacterium tuberculosis drug resistance. Nat Commun. 2026. doi:10.1038/s41467-026-72431-7

30. Wang Q, Boshoff HIM, Harrison JR, Ray PC, Green SR, Wyatt PG, et al. PE/PPE proteins mediate nutrient transport across the outer membrane of. Science. 2020;367: 1147–1151.

31. Gan M, Wang D, Li S, Wang Q, Liu Q. Ongoing evolution of PE/PPE genes in associated with drug resistance and host immune response. mSystems. 2025;10: e0089825.

32. Holt KE, McAdam P, Thai PVK, Thuong NTT, Ha DTM, Lan NN, et al. Frequent transmission of the Mycobacterium tuberculosis Beijing lineage and positive selection for the EsxW Beijing variant in Vietnam. Nat Genet. 2018;50: 849–856.

33. Huang C-C, Trevisi L, Becerra MC, Calderón RI, Contreras CC, Jimenez J, et al. Spatial scale of tuberculosis transmission in Lima, Peru. Proc Natl Acad Sci U S A. 2022;119: e2207022119.

34. Phelan J, O’Sullivan DM, Machado D, Ramos J, Whale AS, O’Grady J, et al. The variability and reproducibility of whole genome sequencing technology for detecting resistance to anti-tuberculous drugs. Genome Med. 2016;8: 132.

35. Marin M, Vargas R, Harris M, Jeffrey B, Epperson LE, Durbin D, et al. Benchmarking the empirical accuracy of short-read sequencing across the M. tuberculosis genome. Bioinformatics. 2022;38: 1781–1787.

36. Godfroid M, Dagan T, Merker M, Kohl TA, Diel R, Maurer FP, et al. Insertion and deletion evolution reflects antibiotics selection pressure in a Mycobacterium tuberculosis outbreak. PLoS Pathog. 2020;16: e1008357.

37. Gagneux S, DeRiemer K, Van T, Kato-Maeda M, de Jong BC, Narayanan S, et al. Variable host-pathogen compatibility in Mycobacterium tuberculosis. Proc Natl Acad Sci U S A. 2006;103: 2869–2873.

38. Cohen KA, Bishai WR, Pym AS. Molecular Basis of Drug Resistance in Mycobacterium tuberculosis. Microbiol Spectr. 2014;2. doi:10.1128/microbiolspec.MGM2-0036-2013

39. Mortimer TD, Weber AM, Pepperell CS. Signatures of Selection at Drug Resistance Loci in. mSystems. 2018;3. doi:10.1128/mSystems.00108-17

40. Houghton J, Rodgers A, Rose G, D’Halluin A, Kipkorir T, Barker D, et al. The Mycobacterium tuberculosis sRNA F6 Modifies Expression of Essential Chaperonins, GroEL2 and GroES. Microbiol Spectr. 2021;9: e0109521.

41. Bar-Oz M, Martini MC, Alonso MN, Meir M, Lore NI, Miotto P, et al. The small non-coding RNA B11 regulates multiple facets of Mycobacterium abscessus virulence. PLoS Pathog. 2023;19: e1011575.

42. Wan L, Hu P, Zhang L, Wang Z-X, Fleming J, Ni B, et al. Omics analysis of Mycobacterium tuberculosis isolates uncovers Rv3094c, an ethionamide metabolism-associated gene. Commun Biol. 2023;6: 156.

43. Dupuy P, Ghosh S, Adefisayo O, Buglino J, Shuman S, Glickman MS. Distinctive roles of translesion polymerases DinB1 and DnaE2 in diversification of the mycobacterial genome through substitution and frameshift mutagenesis. Nat Commun. 2022;13: 4493.

44. Mycobrowser. [cited 6 May 2026]. Available: https://mycobrowser.epfl.ch/

45. Saund K, Snitkin ES. Hogwash: three methods for genome-wide association studies in bacteria. Microb Genom. 2020;6. doi:10.1099/mgen.0.000469

46. Murphy KC, Nelson SJ, Nambi S, Papavinasasundaram K, Baer CE, Sassetti CM. ORBIT: a New Paradigm for Genetic Engineering of Mycobacterial Chromosomes. mBio. 2018;9. doi:10.1128/mBio.01467-18

47. Sysoeva TA, Zepeda-Rivera MA, Huppert LA, Burton BM. Dimer recognition and secretion by the ESX secretion system in Bacillus subtilis. Proc Natl Acad Sci U S A. 2014;111: 7653–7658.

48. Poulsen C, Panjikar S, Holton SJ, Wilmanns M, Song Y-H. WXG100 protein superfamily consists of three subfamilies and exhibits an α-helical C-terminal conserved residue pattern. PLoS One. 2014;9: e89313.

49. Solomonson M, Setiaputra D, Makepeace KAT, Lameignere E, Petrotchenko EV, Conrady DG, et al. Structure of EspB from the ESX-1 type VII secretion system and insights into its export mechanism. Structure. 2015;23: 571–583.

50. Abramson J, Adler J, Dunger J, Evans R, Green T, Pritzel A, et al. Accurate structure prediction of biomolecular interactions with AlphaFold 3. Nature. 2024;630: 493–500.

51. Sani M, Houben ENG, Geurtsen J, Pierson J, de Punder K, van Zon M, et al. Direct visualization by cryo-EM of the mycobacterial capsular layer: a labile structure containing ESX-1-secreted proteins. PLoS Pathog. 2010;6: e1000794.

52. Mesman AW, Baek S-H, Huang C-C, Kim Y-M, Cho S-N, Ioerger TR, et al. Characterization of Drug-Resistant Lipid-Dependent Differentially Detectable. J Clin Med. 2021;10. doi:10.3390/jcm10153249

53. Quispe N, Asencios L, Obregon C, Velásquez GE, Mitnick CD, Lindeborg M, et al. The fourth national anti-tuberculosis drug resistance survey in Peru. Int J Tuberc Lung Dis. 2020;24: 207–213.

54. Lim ZL, Drever K, Dhar N, Cole ST, Chen JM. Mycobacterium tuberculosis EspK has active but distinct roles in the secretion of EsxA and EspB. J Bacteriol. 2022;204: e0006022.

55. Zhang H, Medina-Jaudes N, Forcada-Nadal A, Harrison EM, Coll F. In host mutational adaptation of Mycobacterium tuberculosis complex strains during tuberculosis infection. J Infect Dis. 2026. doi:10.1093/infdis/jiag209

56. Song H, Sandie R, Wang Y, Andrade-Navarro MA, Niederweis M. Identification of outer membrane proteins of Mycobacterium tuberculosis. Tuberculosis (Edinb). 2008;88: 526–544.

57. Sundar S, Thangamani L, Piramanayagam S. Computational identification of significant immunogenic epitopes of the putative outer membrane proteins from Mycobacterium tuberculosis. J Genet Eng Biotechnol. 2021;19: 48.

58. Sidders B, Pirson C, Hogarth PJ, Hewinson RG, Stoker NG, Vordermeier HM, et al. Screening of highly expressed mycobacterial genes identifies Rv3615c as a useful differential diagnostic antigen for the Mycobacterium tuberculosis complex. Infect Immun. 2008;76: 3932–3939.

59. Cole ST, Brosch R, Parkhill J, Garnier T, Churcher C, Harris D, et al. Deciphering the biology of Mycobacterium tuberculosis from the complete genome sequence. Nature. 1998;393: 537–544.

60. Mitra A, Ko Y-H, Cingolani G, Niederweis M. Heme and hemoglobin utilization by Mycobacterium tuberculosis. Nat Commun. 2019;10: 4260.

61. Singh P, Kaufman CB, Whitworth L, Stubbendieck RM, Morgenstein R, Wozniak KL, et al. PPE64 is a mycomembrane channel protein that functions in heme iron uptake and moonlights in biofilm formation in. mBio. 2026;17: e0328125.

62. Su H, Zhang Z, Liu Z, Peng B, Kong C, Wang H, et al. PPE60 antigen drives Th1/Th17 responses via Toll-like receptor 2-dependent maturation of dendritic cells. J Biol Chem. 2018;293: 10287–10302.

63. Mirkin SM. Expandable DNA repeats and human disease. Nature. 2007;447: 932– 940.

64. Liu Q, Liu YJ, Liu R, Culviner PH, Wang X, Wolf ID, et al. High-throughput cytological profiling uncovers genotype-phenotype associations in Mycobacterium tuberculosis clinical isolates. mSystems. 2025;10: e0097225.

65. Aguilar-Ayala DA, Tilleman L, Van Nieuwerburgh F, Deforce D, Palomino JC, Vandamme P, et al. The transcriptome of Mycobacterium tuberculosis in a lipid-rich dormancy model through RNAseq analysis. Sci Rep. 2017;7: 17665.

66. Danilchanka O, Mailaender C, Niederweis M. Identification of a novel multidrug efflux pump of Mycobacterium tuberculosis. Antimicrob Agents Chemother. 2008;52: 2503–2511.

67. Mehta PK, Pandey AK, Subbian S, El-Etr SH, Cirillo SLG, Samrakandi MM, et al. Identification of Mycobacterium marinum macrophage infection mutants. Microb Pathog. 2006;40: 139–151.

68. Ruley KM, Ansede JH, Pritchett CL, Talaat AM, Reimschuessel R, Trucksis M. Identification of Mycobacterium marinum virulence genes using signature-tagged mutagenesis and the goldfish model of mycobacterial pathogenesis. FEMS Microbiol Lett. 2004;232: 75–81.

69. Houben ENG, Bestebroer J, Ummels R, Wilson L, Piersma SR, Jiménez CR, et al. Composition of the type VII secretion system membrane complex. Mol Microbiol. 2012;86: 472–484.

70. Bottai D, Di Luca M, Majlessi L, Frigui W, Simeone R, Sayes F, et al. Disruption of the ESX-5 system of Mycobacterium tuberculosis causes loss of PPE protein secretion, reduction of cell wall integrity and strong attenuation. Mol Microbiol. 2012;83: 1195–1209.

71. Abdallah AM, Verboom T, Weerdenburg EM, Gey van Pittius NC, Mahasha PW, Jiménez C, et al. PPE and PE_PGRS proteins of Mycobacterium marinum are transported via the type VII secretion system ESX-5. Mol Microbiol. 2009;73: 329– 340.

72. Pallen MJ. The ESAT-6/WXG100 superfamily -- and a new Gram-positive secretion system? Trends Microbiol. 2002;10: 209–212.

73. Chen X, Cheng H-F, Zhou J, Chan C-Y, Lau K-F, Tsui SK-W, et al. Structural basis of the PE-PPE protein interaction in. J Biol Chem. 2017;292: 16880–16890.

74. Comas I, Coscolla M, Luo T, Borrell S, Holt KE, Kato-Maeda M, et al. Out-of-Africa migration and Neolithic coexpansion of Mycobacterium tuberculosis with modern humans. Nat Genet. 2013;45: 1176–1182.

75. Langmead B, Salzberg SL. Fast gapped-read alignment with Bowtie 2. Nat Methods. 2012;9: 357–359.

76. Benson G. Tandem repeats finder: a program to analyze DNA sequences. Nucleic Acids Res. 1999;27: 573–580.

77. Cingolani P. Variant Annotation and Functional Prediction: SnpEff. Methods Mol Biol. 2022;2493: 289–314.

78. Stamatakis A. RAxML version 8: a tool for phylogenetic analysis and post-analysis of large phylogenies. Bioinformatics. 2014;30: 1312–1313.

79. Gutierrez MC, Brisse S, Brosch R, Fabre M, Omaïs B, Marmiesse M, et al. Ancient origin and gene mosaicism of the progenitor of Mycobacterium tuberculosis. PLoS Pathog. 2005;1: e5.

80. Mulholland CV, Wiggins TJ, Cui J, Vilchèze C, Rajagopalan S, Shultis MW, et al. Propionate prevents loss of the PDIM virulence lipid in Mycobacterium tuberculosis. Nat Microbiol. 2024;9: 1607–1618.

81. Mehra A, Philips JA. Analysis of Mycobacterial Protein Secretion. Bio Protoc. 2014;4. doi:10.21769/bioprotoc.1159

82. Oluoch PO, Koh E-I, Proulx MK, Reames CJ, Papavinasasundaram KG, Murphy KC, et al. Chemical genetic interactions elucidate pathways controlling tuberculosis antibiotic efficacy during infection. Proc Natl Acad Sci U S A. 2025;122: e2417525122.

